# Acclimatization potential in stress tolerant Caribbean corals - outcomes of a 5 year cross-shelf reciprocal transplant experiment

**DOI:** 10.64898/2026.09.26.754506

**Authors:** Calista L. Hundley, Lily Kratz, Colleen B. Bove, Lisa Carne, Justin H. Baumann

## Abstract

The identification and use of stress-tolerant coral in outplanting efforts has been proposed as a method of restoring and bolstering reef health under increasing environmental change. Prior research has focused on understanding survival in cross-reef transplantation, but evidence of long term resiliency of outplanted corals remains limited. Here, we report data at a five-year timepoint of a reciprocal transplant study of the stress-tolerant Caribbean corals *Pseudodiploria strigosa* and *Siderastrea siderea* between a nearshore and offshore site in Belize. Extreme marine heat wave conditions in August of 2024 revealed species and site-specific bleaching patterns. Through assessments of mortality, energy reserves, and chlorophyll concentrations, *P. strigosa* appear to be more limited in long-term acclimatization potential to the nearshore environment while *S. siderea* maintain acclimatization capacity displayed 17 months post transplant to nearshore. This work conveys information relevant to managers in selecting species for restoration efforts based on short-term growth and long-term resilience in native versus non-native sites, offers insight into potential mechanisms driving these trends, and takeaways on site specific management decisions for successful coral outplanting success.

## Introduction

Coral reefs are some of the most biodiverse and valuable ecosystems on the planet with more than half a billion people globally relying on reef ecosystems for coastal protection, food, and to support their livelihoods (Grigg et al. 1984; Wilkinson 2004). The economic value associated with tourism alone on coral reefs is estimated to generate $35.8 billion (USD) a year globally (Spalding et al. 2017). However, environmental conditions on coral reefs are dramatically shifting due to global change stressors, including more frequent and intense marine heat wave events (Oliver et al. 2018). These extreme marine heat events result in increased coral mortality that ultimately leads to widespread coral reef decline (Leggat et al. 2019). Simultaneously, coral reefs are increasingly threatened by direct human impacts, such as overfishing, sedimentation, and pollution (Wilkinson 2004). Together, these direct and indirect challenges are further exacerbated at local scales, including in the Caribbean Sea and Mesoamerican Barrier Reef System (MBRS) where more than 75% of the reefs are considered threatened (Burke et al. 2011), resulting in a loss of 48% of hard coral cover from 1980 to 2024 (Green et al. 2008; Alves et al. 2022; Wicquart et al. 2025).

While coral reefs face unprecedented rates of decline, evidence suggests that natural variation in environmental tolerance among coral genotypes may offer a path forward for reef persistence and restoration success. For example, phenotypic plasticity, or the ability of an organism to display altered phenotypes in response to environmental variation, may act as a mechanism of acclimatization to support persistence of reef-building corals on future reefs (Seebacher et al. 2015; Liew et al. 2020). For instance, species of *Porites* corals in Moorea, French Polynesia displayed the potential to acclimatize to changes in turbidity within weeks when transplanted between different reef sites (Padilla-Gamiño et al. 2012). Other acclimatization mechanisms in corals include epigenetic gene modification, which can occur quickly in response to environmental cues without modification of the underlying genotype (Hackerott et al. 2023), and increased production of heat shock proteins and antioxidants (Liew et al. 2020).

Beyond these physiological mechanisms for acclimatization, a corals’ prior environmental history can further modulate its stress response, with exposure to sublethal levels of stress known to prepare corals to better face future environmental challenges. Pooled data from across five global coral reefs suggests that daily thermal variability is the most influential predictor of coral bleaching, with higher thermal variability associated with reduced odds of massive bleaching events (Safaie et al. 2018). Daily thermal variability can also modulate photophysiology, with exposure to high variability treatments resulting in less negative impacts of heat stress on Fv/Fm in *Porites rus* and *Acropora hyacinthus* (Cheh 2025). Further, repeated sublethal high temperature events led to induced increased thermal tolerance for corals on the Great Barrier Reef (Ainsworth et al. 2016). While such environmental priming is an interesting acclimatory pathway, it may only occur on small spatial scales and on reefs that are not subject to increasingly lethal marine heatwaves as global temperatures continue to increase (Ainsworth et al. 2016). Therefore, considerations of how prior environmental history may impact potential for acclimatization in tropical corals, as well as the limits of the mechanisms driving them are important in the maintenance of coral species under increasingly stressful environments.

While acclimatization mechanisms are critical for short-term coral survival, their effectiveness is inherently constrained by a species’ thermal history. Specifically, species adapted to thermally extreme environments like the Caribbean typically display restricted thermal tolerance windows and are predicted to be at a disadvantage under additional increases in global temperatures (Somero 2010). It is theorized that organisms adapted to survive in thermally extreme environments (*i.e.,* the tropics) may be more limited in ability to acclimatize to novel thermal extremes (Stillman 2003). Specifically, constraints to acclimatization may be due to the complex partnerships of the coral host with their various symbionts (*e.g.*, eukaryotic endosymbiotic microalgae, and prokaryotic associates) referred to collectively as the coral holobiont (Voolstra et al. 2021; Bove et al. 2022b). Previous studies spanning Pacific reef building coral and temperate corals show evidence of symbiont plasticity as a major driver in holobiont acclimatization when exposed to new thermal environments, pointing to potentially conserved limits in host plasticity as compared to symbiont role (Howells et al. 2013; Rodolfo-Metalpa et al. 2014). Understanding the acclimatization potential of corals along with the mechanisms supporting such acclimatization, provides important insight into coral capacity to withstand future stress.

In the Caribbean, *Siderastrea siderea* and *Pseudodiploria strigosa* represent two historically common and stress-tolerant reef-building coral species that offer valuable models for examining acclimatization potential under increasingly challenging environmental conditions (Darling et al. 2012). Despite their hardiness and prevalence on reefs, these stress-tolerant coral species have not been the target in Caribbean coral restoration programs due to their relatively slow growth rates compared to faster growing corals, such as *Acropora sp*. In Caribbean reef restoration, many projects have emphasized the benefit of using *Acropora* species (principally *Acropora cervicornis*) due to fast growth rates, morphology, physiological plasticity, and recruitment success in laboratory settings which supports fragmentation and outplanting efforts (Hodges and Hallock 2025). Additionally, the branching morphology of *Acropora sp.* results in more rapid recovery of reef function due to improved three-dimensional complexity, which provides shelter for fishes and invertebrates and helps protect islands and coastline from erosion by dissipating wave energy (Harris et al. 2018). However, stress-tolerant corals are usually more thermally tolerant than these competitive fast-growing corals and therefore more frequently make up a sizable amount of modern living reefs given their relative success surviving extreme heat events like heatwaves (Sommer et al. 2014; Toth et al. 2019; Alves et al. 2022). Researchers have since proposed proactive selection of heat-tolerant genera of corals as one recommendation for restoration planners to enhance restoration efforts (Caruso et al. 2021). Additionally, coral genotypes that display higher stressor-specific tolerance are increasingly employed, especially when the goal is to restore more sensitive species (Humanes et al. 2024; Klepac et al. 2024). In general the use of genetically diverse coral fragments provides greater opportunity for success in restoration efforts against factors like environmental stress or disease presence (Hodges and Hallock 2025). Restoration programs are growing in size and scope as practitioners are investing in adding more species and genetic diversity to their restoration plans (Cabaitan et al. 2015; O’Donnell et al. 2018). Simultaneously, careful site selection and documentation of shared traits that predict success are important considerations as practitioners work against a warming ocean, local stressors, and novel diseases to preserve biodiversity and reef functionality (Schill et al. 2021; Yuen et al. 2023).

Here, we employ a nearshore to offshore reciprocal-transplant experiment to assess the acclimatization potential of *S. siderea* and *P. strigosa* from the southern portion of the Belize MBRS. Reciprocal transplant experiments can be an effective method to differentiate signals of genetic adaptation and phenotypic plasticity that inform a given coral’s response to stress (Kawecki and Ebert 2004; Palumbi et al. 2014). We report the physiological responses of both species 5 years post transplantation across treatments. These results build on previous findings reported over the first year and a half of the transplant experiment (Baumann et al. 2021). Our results provide important context to coral resilience and restoration practices by extending monitoring and reporting on transplantation success beyond the typical timeframe (Boström-Einarsson et al. 2020). Further, this work has the potential to contribute to site and species selection for coral restoration efforts in the region and can shed light on the complex nature of resilience in Caribbean corals.

## 2 Methods

### 2.1 Experimental Design

The reciprocal transplant experiment is outlined in detail in Baumann et al. (2021). Briefly, transplant sites were selected given their geographical separation and differing environmental regimes (Baumann et al. 2016). False Caye is located in the nearshore lagoon ∼2km from the mainland (16.602554°N, 88.340223°W), while Silk Caye is located about 30 km offshore in the backreef of the Belize Mesoamerican Barrier Reef System (16.45026°N, 88.04360°W). In the Belize Mesoamerican Barrier Reef System, False Caye and other nearshore reefs have higher nutrient and sediment loads and are more prone to extreme temperatures compared to offshore reefs (Baumann et al. 2016). As previously reported by Baumann et al. (2021), the mean *in situ* water temperatures differed between the sites over the first 17-months of the transplant period, where False Caye reef nearshore (NS) was 0.4 ± 0.13°C warmer than the offshore Silk Caye reef (OS). Finally, the tables holding the transplanted corals were located at different depths at each (NS table at ∼1.22 meters, OS table at ∼1.83 meters), suggesting differences in light exposure.

At the start of this transplantation study (December 2017), both the NS and OS site contained six parent colonies – assumed to be genetically unique genotypes – of both coral species (Baumann et al. 2021). Each parent colony was cut into 13 smaller fragments using a wet tile saw; 1 fragment was frozen as a control at the start of the experiment, 6 fragments were placed in the NS and 6 fragments were placed in the OS. For our analysis, fragments placed at the same site as their parent colony are referred to as natives (*i.e.*, “NS Native” and “OS Native”) and those placed at the opposite site are referred to as the transplants (*i.e.*, “Transplant to NS” and “Transplant to OS”). Fragments were assessed for mortality, bleaching, buoyant weight, and preserved for downstream physiological assays at the start of the experiment (December 2017), three (March 2018), nine (October 2019), and 17 months (May 2019) post transplant.

Sixty months (5 years) after the start of the experiment, a total of 68 surviving fragments remained across the two sites: 45 NS (24 *S. siderea* and 21 *P. strigosa*) and 23 OS (12 *S. siderea* and 11 *P. strigosa*). Algae and other fouling organisms were removed from fragments before buoyantly weighing each in triplicate to calculate % change in buoyant weight over the course of the experiment (Jokiel et al. 1978). Photosynthetic efficiency of photosystem II (F_v_/F_m_) was measured for each fragment using a diving pulse amplitude-modulated (PAM) Fluorometer (Walz, Germany)(Beer et al. 1998). After 30-minutes of dark acclimation to ensure accurate fluorometry readings, triplicate measurements were taken by random placement of the photon probe on each coral fragment (Fitt et al. 2001; Warner et al. 2010). A subset of these fragments was preserved by flash freezing in liquid nitrogen for physiological analyses (n = 7-9 per species per treatment, 34 total) with a goal of collecting at least one individual from each original parent colony. Flash-frozen fragments were transported back to Mount Holyoke College (South Hadley MA, USA) where they were stored at -80°C for downstream analysis.

### 2.2 Identification of coral fragments with degraded labels

Some corals that remained on the experimental tables in August 2023 were unidentifiable due to the degradation or loss of labels (n = 13, 4 *P. strigosa*, 9 *S. siderea*). In an effort to match these unknown corals with their original parent colony for downstream analyses, we collected small tissue samples from all corals for 2b-RADseq analysis. DNA was extracted using a modified phenol-chloroform method (Davies et al. 2013) from the 13 unknown samples along with known fragments of each of the 12 original coral colonies per species (n = 25 total samples). The extracted DNA was cleaned using Zymo Genomic RNAse A and DNA Clean and Concentrator kits, concentrations estimated using a SpectraMax QuickDrop, and visualized on an agarose gel. Successfully extracted DNA samples were sent to CD Genomics for 2b-RAD library preparation and sequencing across one lane of Illumina HiSeq X Ten using paired-end 150-base pair (bp) sequencing. The resulting raw data were processed using CD Genomics’ in-house pipeline that included merging paired-end reads, trimming adaptor sequences, removal of terminal 3-bp positions, and filtering out low-quality reads (Zhang et al. 2014).

Cleaned reads were analysed following the 2b-RADseq pipeline presented at https://github.com/z0on/2bRAD_denovo and de novo genome references for both S. siderea and P. strigosa were created following Rippe et al. (2021). To create the de novo references, Symbiodiniaceae contamination was removed by mapping reads from both species to concatenated genomes from four Symbiodiniaceae genera (*Symbiodinium*, *Breviolum*, *Cladocopium*, and *Durusdinium*; (Shoguchi et al. 2013; Aranda et al. 2016; Liu et al. 2018; Dougan 2020) using Bowtie2/2.4.4-rhel8 (Langmead and Salzberg 2012). Samples were then separated by coral species and CD-HIT (4.8.1) was used to cluster and assemble remaining reads into de novo references of 30 pseudochromosomes each (Fu et al. 2012).

The cleaned reads were mapped to the appropriate de novo reference (*S. siderea* or *P. strigosa*) using Bowtie2/2.4.4-rhel8 followed by genotyping and single-nucleotide polymorphism (SNP) identification was carried out using ANGSD/0.934 (Korneliussen et al. 2014). Finally, *hclust* was employed in RStudio/2026.09.0+174 to construct a hierarchical tree based on pairwise identity-by-state values across all samples per species to determine unknown coral genotypes (R Core Team 2026). From this analysis, three *S. siderea* corals with lost ID tags were able to be confirmed to the colony and treatment level: one fragment from OS colony six that was transplanted to NS and two NS native fragments from colonies two and six, respectively. Four *P. strigosa* and six *S. siderea* fragments that survived through the 60 month timepoint remained unidentified (Tables S2, S3).

### 2.3 Physiological assays

Preserved coral fragments were sectioned into three to four subfragments using a wet tile saw: one subfragment for energy reserves, one for chlorophyll concentration, and remaining fragments were preserved at -80°C. Subfragments were ground, both tissue and skeleton, into a homogenous paste using a mortar and pestle, decanted into a 50 ml conical tube, and suspended in 5-10 ml of MilliQ (Grottoli et al. 2004). Samples were vortexed and then centrifuged at 4000 rpm for 10 minutes to separate the skeleton from the tissue. The tissue fraction (supernatant) was poured off into a clean 50 ml conical. Tissue slurry was homogenized at ∼28000 rpm for 6 minutes (Polytron 3100). Bulk tissue samples (coral and symbiont material) were used for energy reserve assays while chlorophyll concentration assays required separation of symbiont tissue from the coral tissue via centrifugation (Baumann et al. 2021).

#### Energy Reserve Analysis

After homogenization, a 1 mL aliquot of bulk tissue was prepared for each individual biological macromolecule assay of protein, lipid, and carbohydrate. Protein concentrations were assessed using a modified protocol based on the Bradford colorimetric assay method where samples prepared with Coomassie Blue Dye were run alongside Bovine serum albumin based standards in a microplate reader at 595 nm (BioTek Synergy H1 Hybrid Reader)(Bradford 1976; Mclachlan et al. 2020; Baumann et al. 2021). Total lipids were extracted using phosphoric-sulfuric acid method and measured alongside corn oil standards in a microplate reader at a wavelength of 540 nm (Bove et al. 2022a). A phenol-sulfuric acid protocol (DuBois et al. 1956) modified for 96-well plates (Masuko et al. 2005; Bove et al. 2022a) was used for carbohydrate extraction and quantification using D-glucose standards in a microplate reader at a wavelength of 485 nm. The absorbance values of the standards were plotted as a standard curve. Mean absorbance from three replicate measurements was compared against standard curves to estimate respective bulk energy reserve concentration (mg/mL) of each coral sample. Bulk energy reserve content (mg) was determined by multiplying the concentration by the original bulk tissue slurry volume (mL) and employed dilution factors and then standardized to total biomass (mg/mg) quantified via ash-free dry weight of tissue (Grottoli et al. 2004).

#### Chlorophyll Assay

Chlorophyll-a and chlorophyll-c2 molecules were extracted from symbiont cells using a modified methanol extraction protocol (Warren 2008). Mean absorbance from triplicate measurements of symbiont samples run alongside standards of water, methanol, and healthy leaf tissue at 632 nm and 665 nm was used to calculate chlorophyll concentrations (μg/cm^3^)(Ritchie 2006). Total chlorophyll molecule contents (μg) were determined by multiplying the concentrations by the original subfragment surface area (cm^3^). Surface area estimation of each subfragment was conducted using planar projection photography and image analysis using ImageJ (Mclachlan and Grottoli 2021).

### 2.4 Statistical Analysis

To assess the effect of transplant treatment on coral physiology (chlorophyll concentrations, photosynthetic efficiency, protein concentrations, carbohydrate concentrations, lipid concentrations), analyses were conducted separately for *P. strigosa* and *S. siderea*. Prior to model fitting, data distributions were visually inspected using histograms, density plots, and quantile-quantile (Q-Q) plots to assess normality.

For both species, we first attempted to fit linear mixed-effects models with native environment, transplant environment, and their interaction as fixed effects, and colony as a random effect to account for non-independence of fragments originating from the same colony (package *lme4*) (Bates et al. 2015). In some cases, the models produced singular fits, indicating that colony-level variance was negligible, and the random effect could not be reliably estimated. In these cases, a simple linear model with treatment as a predictor was used (**see Tables S2**, **S4**).

Note that for *P. strigosa*, one treatment group (Transplant to NS) was excluded from analysis due to insufficient replication (n = 1). Similarly, no *S. siderea* corals survived in the Transplant to OS treatment and thus this treatment level is not present in this species. Model assumptions (homogeneity of variance, normality of residuals) were visually assessed using the *check_model* function (package *performance*)(Lüdecke et al. 2021). Significant treatment effects were followed up with pairwise comparisons using estimated marginal means (package *emmeans*)(Lenth and Piaskowski 2026), with p-values adjusted for multiple comparisons using the Tukey method. All statistical analyses were performed in R (version 4.6.0).

## 3 Results

### Survivorship

Each treatment (NS Native, OS Native, Transplant to NS, Transplant to OS) has an original sample size of 20-22 coral fragments for each species, excluding fragments collected for downstream analysis at each timepoint (T1-3 months, T2 9 months, T3 18 months). After 60 months, NS Native corals had the highest survival rate in *P. strigosa*, with all other treatments producing low survival (<30%; Table 1). NS *P. strigosa* were 56% more likely to survive in their home environment than when transplanted to the OS environment and OS *P. strigosa* were 22% more likely to survive in their home environment than when transplanted to the NS (Table 1). A similar trend was observed in NS *S. siderea*, which were 40% more likely to survive in their home environment than when transplanted to the OS, where they faced 100% mortality (Table 1). Conversely, survival rate of OS Native *S. siderea* was near 50% at home and when transplanted to the nearshore (Table 1).

**Table 1:** Percent survival of corals in each treatment after 60 months. Corals collected at each previous timepoint are omitted from the calculations. Fragments that could not be identified to the treatment level at this timepoint are also omitted (4 *P. strigosa* and 6 *S. siderea*; Table S1). Thus, these numbers are approximate.

| Species | NS native | Transplant to OS | OS Native | Transplant to NS |
| --- | --- | --- | --- | --- |
| *S. siderea* | 40.91% | 0.00% | 55.00% | 50.00% |
| *P. strigosa* | 76.19% | 20.00% | 27.27% | 5.00% |

### Net coral calcification

On average, *S. siderea* fragments grew half as much as *P. strigosa* fragments by 60 months post-transplant (ANOVA, F = 125.49, df = 1, *p* < 0.05; Tukey HSD, *p* < 0.01; **Figure 1**). Transplant to NS *S. siderea* grew by 6.78% ± 1.25%, an amount significantly greater than NS Native (3.47% ± 1.22%; *p* < 0.01) and OS Native *S. siderea* (2.69% ± 1.24%; *p* < 0.01). The elevated growth of Transplant to NS *S. siderea* compared to other treatments mirrors results observed 17 months after transplant. Over the same time frame, NS Native *Pseudodiploria strigosa* grew 8.56% ± 0.63% in 60 months, significantly more than Transplant to OS *P. strigosa* (3.74% ± 1.30%; *p* < 0.01) and OS Native *P. strigosa* (5.15% ± 1.06%; *p* < 0.05).

**Figure 1:**
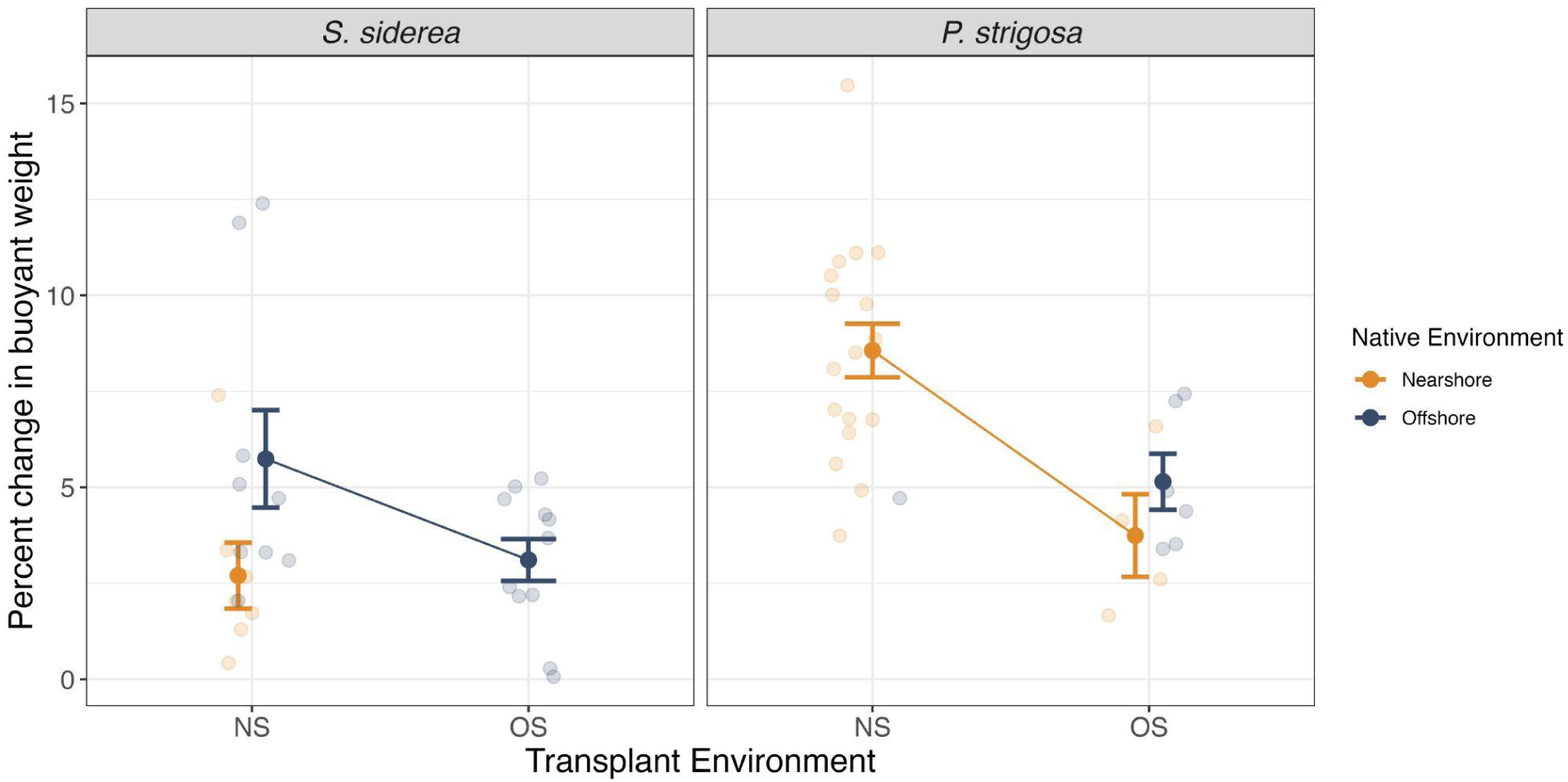
Percent change in skeletal growth. Mean percent change in buoyant weight of *Siderastrea siderea* (left) and *Pseudodiploria strigosa* (right) fragments 60 months post transplantation. Color denotes native collection environment (NS in orange; OS in blue). Raw percent change values are depicted as transparent points with larger, solid points and error bars representing mean ± SE per treatment. No *S. siderea* transplanted to the offshore and only one *P. strigosa* transplanted to the nearshore were alive for analysis, thus these treatment means are not depicted.

### Photosynthetic efficiency

While there was no clear difference in Fv/Fm across treatments in *S. siderea* fragments (F = 1.36, df = 2, *p* = 0.27; **Figure 2**; **Tables S2**, **S3**), treatment did significantly influence Fv/Fm in *P. strigosa* fragments (F = 12.59, df = 2, *p* < 0.001; **Figure 2**; **Table S4**). NS Native *P. strigosa* corals had significantly lower Fv/Fm (mean ± SE = 0.467 ± 0.012) than both OS Native (mean ± SE = 0.550 ± 0.022; Tukey-adjusted *p* < 0.01; **Table S5**) and Transplant to OS corals (mean ± SE = 0.578 ± 0.026; *p* < 0.01; **Table S5**), while corals placed in the OS did not differ significantly from one another, regardless of native environment (*p* = 0.696; **Table S5**).

**Figure 2:**
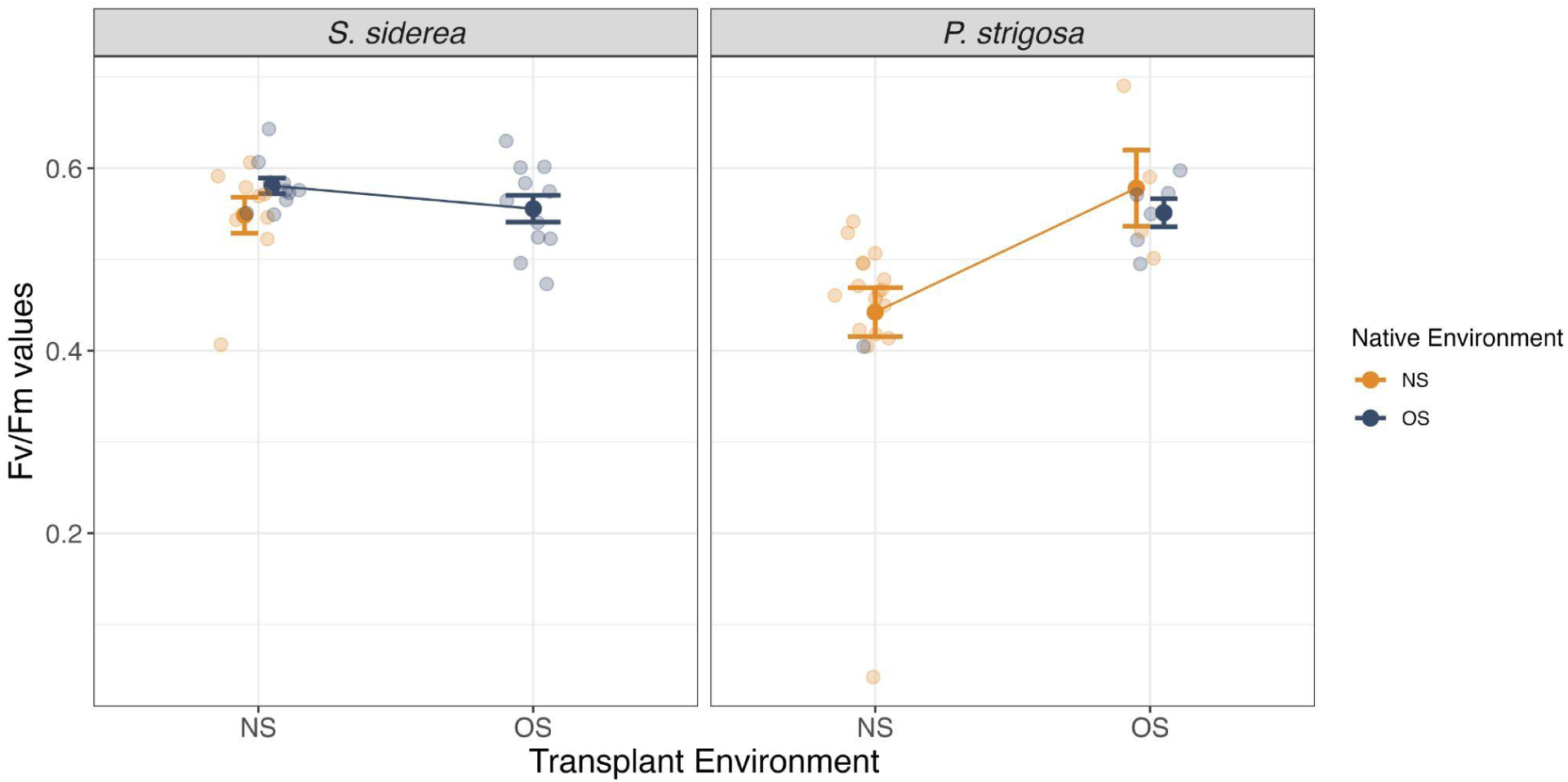
Photosynthetic efficiency 60 months post transplantation. Maximum photosynthetic efficiency (Fv/Fm) of *Siderastrea siderea* (left) and *Pseudodiploria strigosa* (right) fragments. Color denotes native collection environment (NS in orange; OS in blue). Raw fragment Fv/Fm values are depicted as transparent points with larger, solid points and error bars representing mean ± SE per treatment. No *S. siderea* transplanted to the offshore and only one *P. strigosa* transplanted to the nearshore were alive for analysis, thus these treatment means are not depicted.

### Energy reserves

Five years post transplantation, bulk protein concentrations in both *S. siderea* (F = 6.13, df = 2, *p* < 0.05; **Table S2**) and *P. strigosa* (F = 4.47, df = 2, *p* < 0.05; **Table S4**) were significantly impacted by transplant treatment (**Figure 3**). Both NS Native and Transplant to NS corals of *S. siderea* exhibited greater protein concentrations than the OS Native fragments (Tukey-adjusted *p*= 0.044; *p* = 0.048; respectively; **Table S3**). NS Native *P. strigosa* corals had significantly higher protein concentrations than OS Native corals (Tukey-adjusted *p* < 0.05; **Table S5**), while Transplant to OS corals maintained statistically indistinguishable protein concentrations with both NS Native and OS Native corals (*p* = 0.65; *p* = 0.16; respectively; **Table S5**).

**Figure 3:**
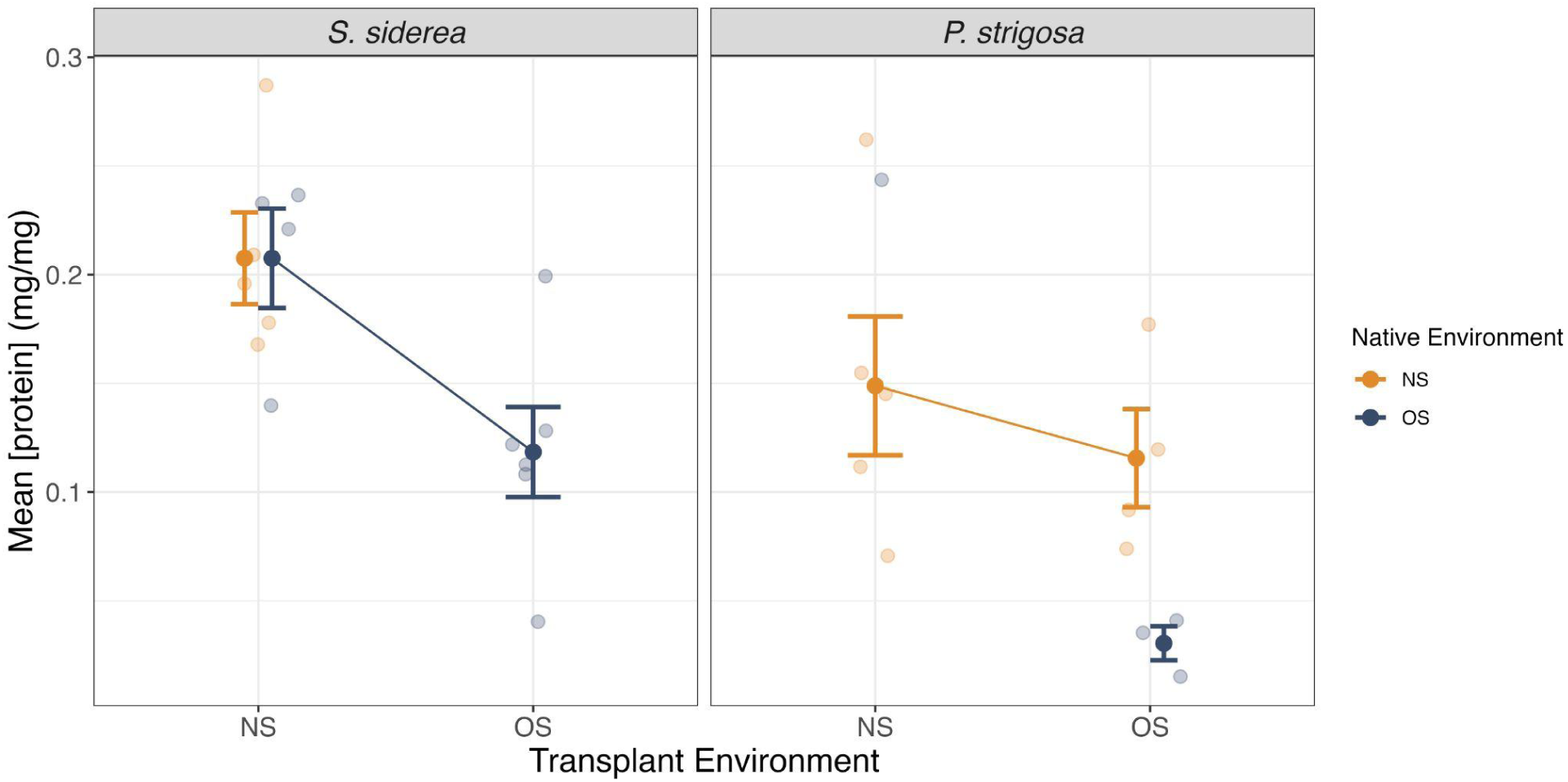
Proteins at 5 year time point. Bulk protein concentrations (mg/mg) of *Siderastrea siderea* (left) and *Pseudodiploria strigosa* (right) fragments 60 months post transplantation. Color denotes native collection environment (NS in orange; OS in blue). No *S. siderea* transplanted to the offshore and only one *P. strigosa* transplanted to the nearshore were alive for analysis, thus these treatment means are not depicted. Raw fragment protein concentrations are depicted as transparent points with larger, solid points and error bars representing mean ± SE per treatment.

*Siderastrea siderea* corals did not exhibit significantly different bulk lipid concentrations regardless of transplant treatment (F = 0.899, df = 2, *p* = 0.43; **Figure 4**; **Table S2**). Similarly, *P. strigosa* lipid concentrations were not statistically different across treatments (F = 1.15, df = 2, *p* = 0.36; **Figure 4**; **Table S4**).

**Figure 4:**
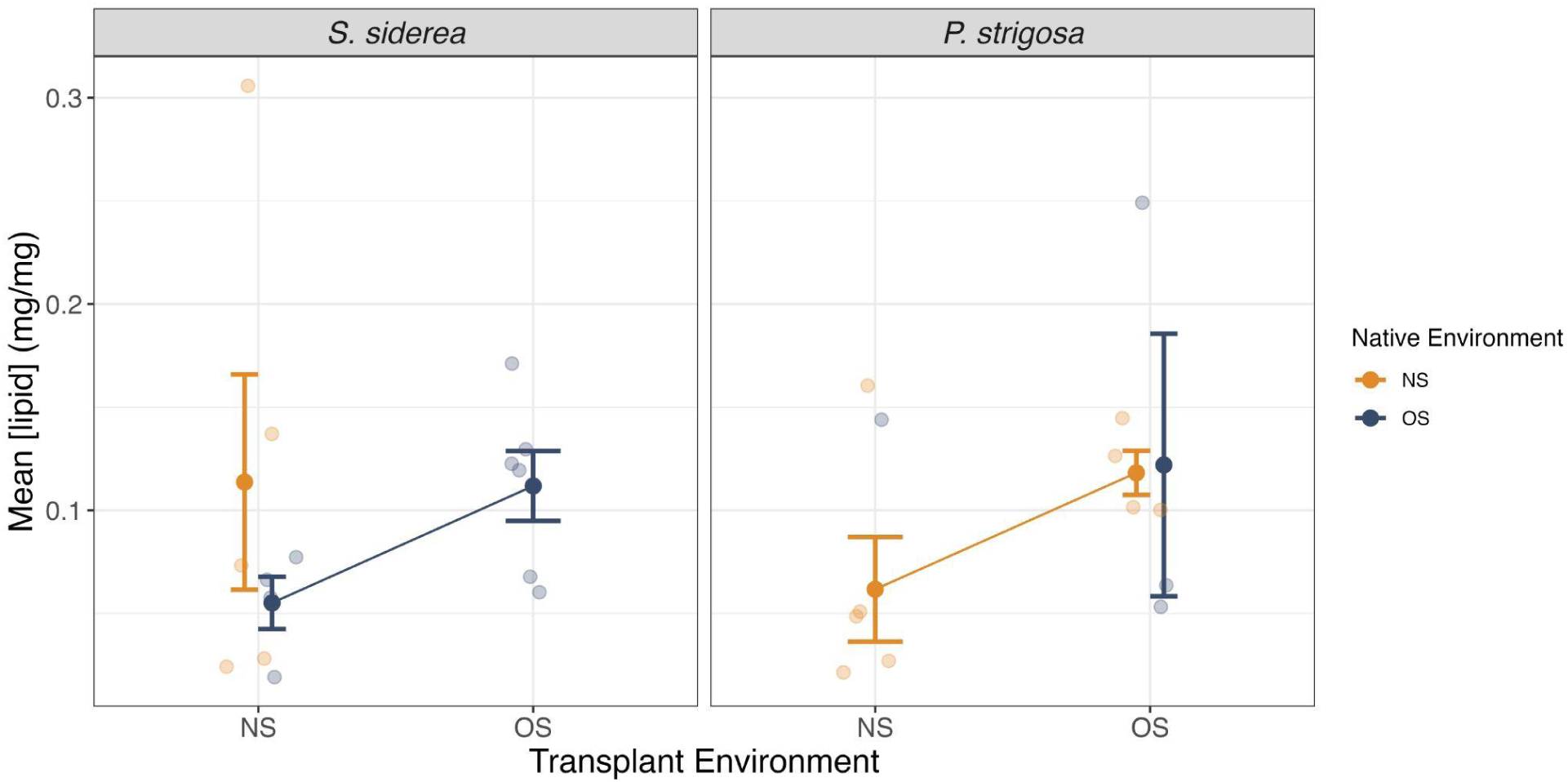
Lipids at 5 year time point. Bulk lipid concentrations (mg/mg) of *Siderastrea siderea* (left) and *Pseudodiploria strigosa* (right) fragments 60 months post transplantation. No *S. siderea* transplanted to the offshore and only one *P. strigosa* transplanted to the nearshore were alive for analysis, thus these treatment means are not depicted. Raw fragment lipid concentrations are depicted as transparent points with larger, solid points and error bars representing mean ± SE per treatment.

Bulk carbohydrate concentrations of *S. siderea* corals were not significantly different from one another across treatments five years post transplant (F = 2.95, df = 2, *p* = 0.09; **Figure 5**; **Table S2**). In *P. strigosa*, however, carbohydrate concentrations differed across treatments (F = 15.22, df = 2, *p* < 0.01; **Figure 5**; **Table S4**), with OS Native corals exhibiting the highest carbohydrate concentrations (Tukey-adjusted *p* < 0.01; **Table S5**) and NS Native and Transplant to OS possessing indistinguishable concentrations from one another (*p* = 0.34; **Table S5**).

**Figure 5:**
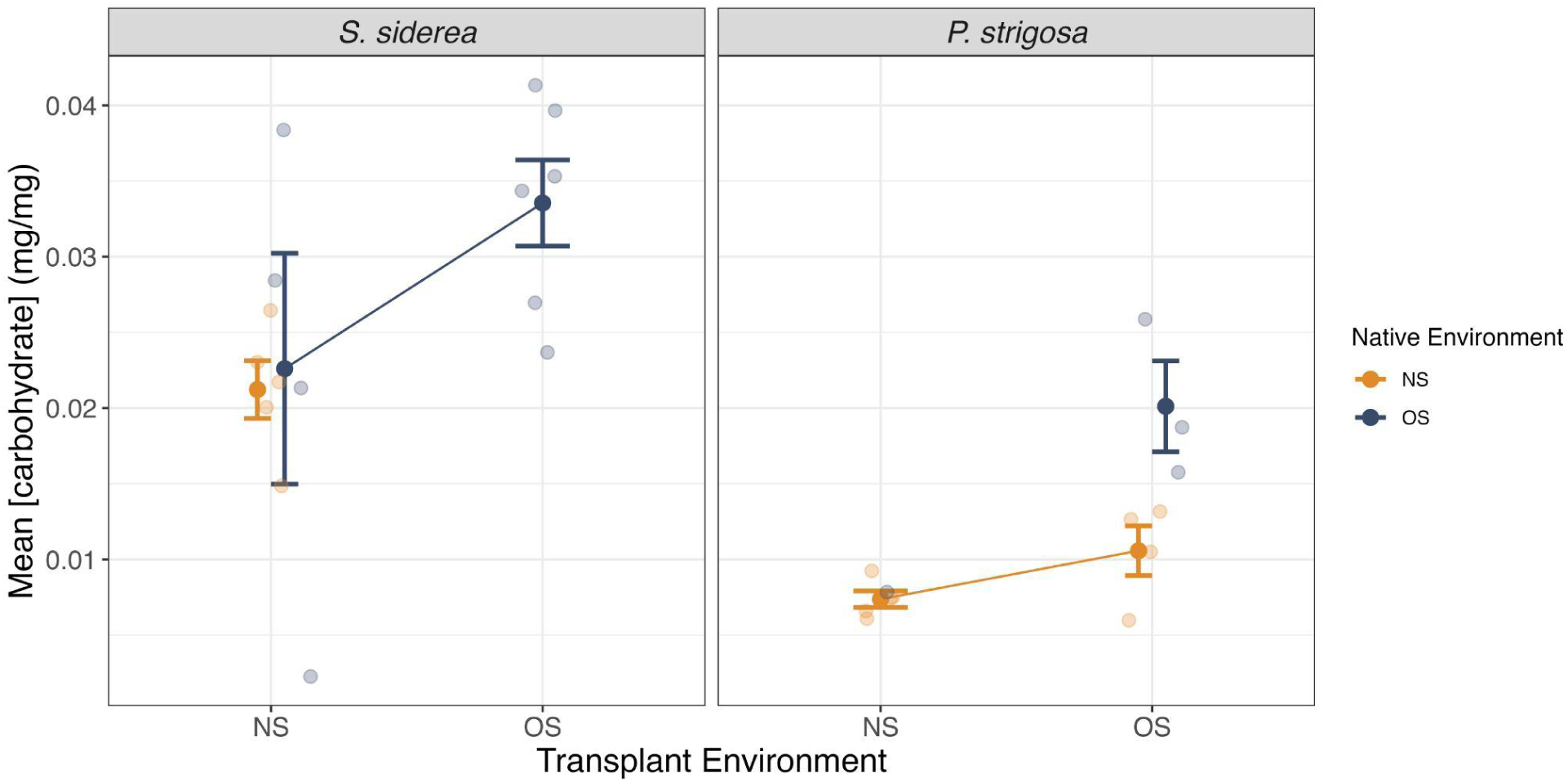
Carbohydrates at 5 year time point. Bulk carbohydrate concentrations (mg/mg) of *Siderastrea siderea* (left) and *Pseudodiploria strigosa* (right) fragments 60 months post transplantation. Color denotes native collection environment (NS in orange; OS in blue). No *S. siderea* transplanted to the offshore and only one *P. strigosa* transplanted to the nearshore were alive for analysis, thus these treatment means are not depicted. Raw fragment carbohydrate concentrations are depicted as transparent points with larger, solid points and error bars representing mean ± SE per treatment.

### Chlorophyll concentrations

Neither chlorophyll-a nor chlorophyll-c2 were significantly different in *S. siderea* fragments across treatments 5 years post transplant (chlorophyll-a: F = 1.86, df = 2, *p* = 0.19; chlorophyll-c2: F = 1.62, df = 2, *p* = 0.24; **Figure 6**; **Table S2**). Conversely, transplantation treatment impacted both chlorophyll pigments in *P. strigosa* corals (chlorophyll-a: F = 21.99, df = 2, *p* < 0.001; chlorophyll-c2: F = 26.11, df = 2, *p* < 0.001; **Figure 6**; **Table S4**). Both pigments were higher in Transplant to OS *P. strigosa* than in both NS Native (chlorophyll-a: *p* < 0.001; chlorophyll-c2: *p* < 0.001; **Table S5**) and OS Native (chlorophyll-a: *p* < 0.01; chlorophyll-c2: *p* < 0.01; **Table S5**) corals.

**Figure 6:**
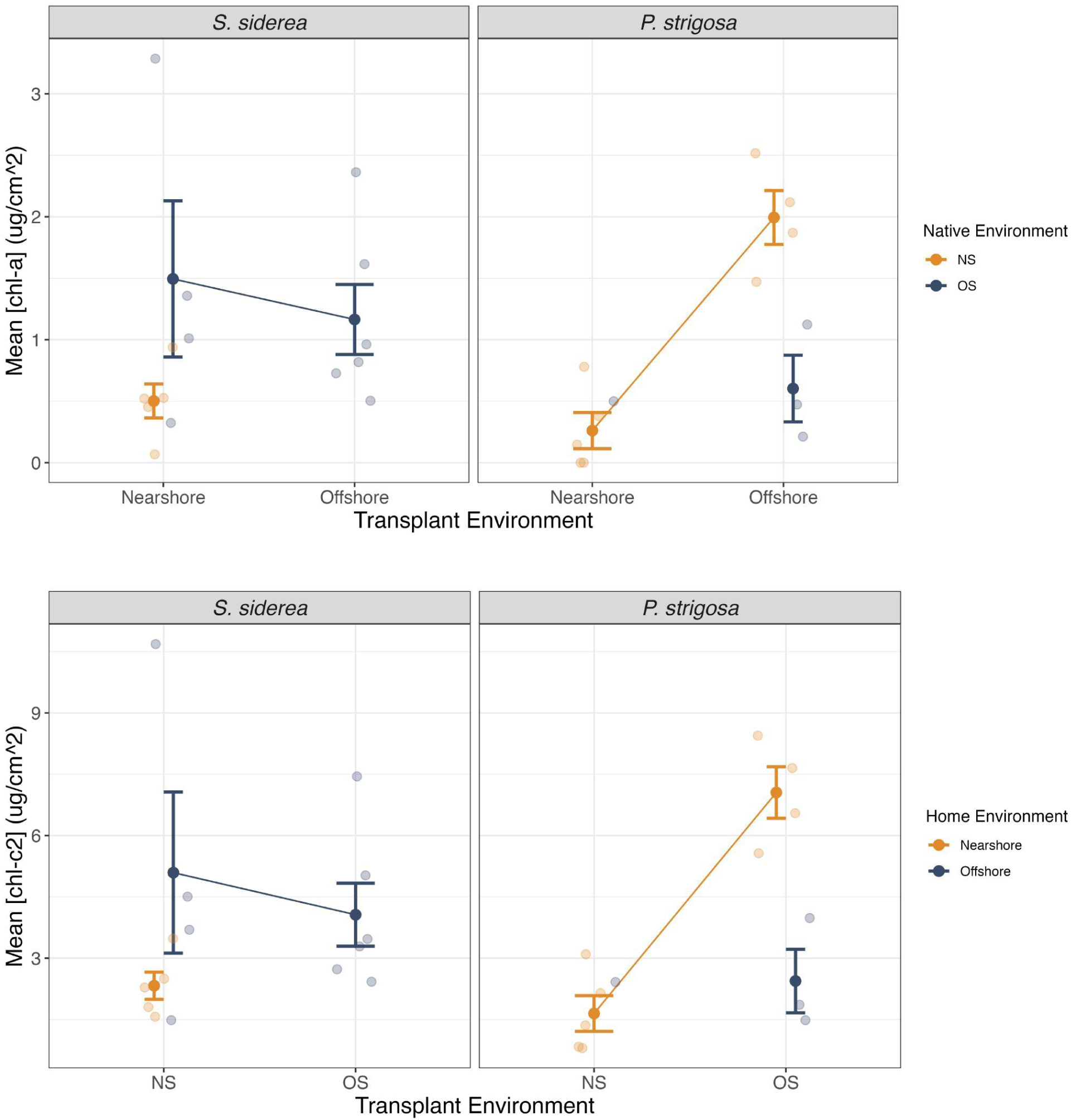
Chlorophyll pigment concentrations at 5 year time point. Chlorophyll-a (top) and chlorophyll-c2 (bottom) concentrations (μg/cm^2^) of *Siderastrea siderea* (left) and *Pseudodiploria strigosa* (right) fragments 60 months post transplantation. Color denotes native collection environment (NS in orange; OS in blue). Raw fragment chlorophyll concentrations are depicted as transparent points with larger, solid points and error bars representing mean ± SE per treatment. No *S. siderea* transplanted to the offshore and only one *P. strigosa* transplanted to the nearshore were alive for analysis, thus these treatment means are not depicted.

## 4 Discussion

We predicted that mortality would be lowest for corals placed in their native environment and highest for corals transplanted to non-native environments due to local adaptation or inability to adjust to new environmental conditions following results from the first 17 months of this study (Baumann et al. 2021). Following these predictions, corals were generally observed to experience less mortality in their native environment by 60 months post transplant. However, *S. siderea* transplanted to the nearshore experienced lower mortality than those native to the nearshore, providing evidence of offshore *S. siderea*’s acclimatization potential. Additionally, *S. siderea* that remained in the offshore (OS Native) had roughly the same survival rate at home vs away (transplanted to NS), whereas transplantation resulted in lower survival in all other scenarios, perhaps due to limited ability to acclimatize to new conditions. Limits in coral acclimatization potential have been proposed due to adaptation of holobionts as well as specific parts of the coral including the endosymbiotic population (Howells et al. 2012, 2013). In terms of adjusting to shifted environmental conditions, the nearshore and offshore sites in this study differ in aspects that could serve as barriers for adjustment. Given the gradient of increasing temperature and variability in the MBRS, the nearshore site would appear as a more thermally extreme site to offshore corals (Baumann et al. 2016). Prolonged exposure to conditions outside of their thermal optimum can result in the mortality of individuals if they cannot adjust (Anttila et al. 2013). Survival through 60 months demonstrates habitat viability for these corals, indicating that it may at least be possible to successfully restore populations of *S. siderea* on degraded nearshore reefs using offshore populations. However, the opposite (seeding the offshore with nearshore genotypes), may not be viable. For *P. strigosa* transplant survival was low in both treatments, indicating that cross-shelf restoration may not be effective on 5-10 years time scales. Notably, it is quite difficult to constrain specific mechanisms of survival and mortality in a field study, especially in an era of frequent heat waves and as a new endemic disease sweeps the region.

Placement of corals in a new environment that is, on average, warmer and contains higher nutrient resources promoting heterotrophic feeding could be beneficial in the short term as it may situate these organisms within a thermal range that moves them closer to their thermal optimum and promotes higher productivity (*i.e.,* growth) (Lough and Cantin 2014; Baumann et al. 2021). The conditions outlined above mirror the transplant to nearshore treatment in our study, where both species exhibited the highest growth rates when transplanted to the nearshore compared to all other treatments (including nearshore native; Baumann et al., 2021). This elevated growth combined with low mortality and was linked with increased heterotrophic opportunity (e.g. more nutrition) using stable isotope analysis (Baumann et al. 2021). *Acropora cervicornis*, another Caribbean coral, was able to mitigate negative effects of temperature stress on growth by increasing heterotrophic feeding rates (Towle et al. 2015). However, in the long term, annual increases in average values and variability of temperature with the shifting climate could become large enough that these thermal values surpass the level that these individuals can survive in, exceeding their thermal optimum (Cooper et al. 2008). This appears to be the case here, as multiple severe marine heatwaves were reported across the 60 month experiment in the nearshore site (Fig S1), including during the 60 month collection and measurement timepoint. Repeated marine heatwaves and bleaching stress may well have pushed the conditions in the nearshore environment from beneficial to stressful for *P. strigosa* corals transplanted to the nearshore, as they suffered considerably more mortality than did their nearshore native conspecifics. However, *S. siderea* corals transplanted to the nearshore survived at near the same rate as they did at home (offshore native) and as conspecifics native to the nearshore, indicating a resilience to this repeated marine heatwave associated stress. Additionally, *S. siderea* transplanted to the nearshore continued to have the highest growth rate after 60 months, mirroring results from the first 17 months of this experiment. Thus, *S. siderea* appear more robust to the stressors facing a nearshore site and appear to maintain their ability to acclimatize to such conditions across 60 months while *P. strigosa* see short term success but were not robust enough to maintain their population after 60 months, likely due to accumulation of stress from repeated heatwaves.

Acclimatization potential of nearshore populations appears to be limited when transplanted to the offshore as net calcification in nearshore native *P. strigosa* was higher than when transplanted to offshore and there were no surviving *S. siderea* in the transplant to offshore treatment. Though mortality and degradation of labels prohibited two-way reaction norms for a full reciprocal design, these results, taken together, support the findings of the short term experiment - that acclimatization / plasticity in populations of both species is limited when transplanted from nearshore to offshore and that both species appear to have elevated growth rates in the nearshore when compared to the offshore.

The appearance of a massive summer heatwave event in 2023 led to record high sea surface temperatures and high mortality across Caribbean reefs (Birkart and Álvarez-Filip 2025). The transplant sites had experienced a cumulative 16 degree heating weeks (*i.e.,* number of weeks the temperature had been 1°C over the bleaching threshold) before corals were collected for analysis of the 60-month timepoint (NOAA Coral Reef Watch 2024). This statistic indicates that widespread bleaching was occurring along the MBRS at the time of collection, as corroborated by our visual surveys, photosynthetic efficiency measurements (F_v_/F_m_), and chlorophyll concentrations of our surviving coral fragments. In addition, the overall lower values of F_v_/F_m_ in *P. strigosa* and the appearance of significantly higher chlorophyll concentrations in nearshore *P. strigosa* transplanted to the offshore support that this species has a lower tolerance and longer recovery from bleaching than *S. siderea* as supported by previous literature (Banks and Foster 2017) and that the offshore site was not as significantly impacted by bleaching as the nearshore site.

Despite their stress-tolerant nature (Darling et al. 2012), *P. strigosa* and *S. siderea* still exhibited declines in physiology under elevated temperature as seen in prior work (Bove et al. 2022a). The energy reserve concentrations of these corals were representative of heavily bleached coral, at reduced values due to the reliance of corals on these reserves in the absence of photoautotrophic or heterotrophic produced energy during bleaching (Grottoli et al. 2006). Both *S. siderea* and *P. strigosa* have been shown to consume particulate organic carbon heterotrophically, possibly underlying their stress tolerance through a nutritional advantage over their less heterotrophic counterparts during stress events, such as bleaching (Mills and Sebens 2004; Mills et al. 2004). It is well known that it can take time for corals to recover their energy reserves post-bleaching and that this recovery period differs by species (Rodrigues and Grottoli 2007; Baumann et al. 2014). There is also a generalized pattern that the recovery of energy reserves follows the successful recovery of symbiont populations and chlorophyll-a pigment (Fitt et al. 2000). If these corals can fully recover their concentrations of energy reserves, they would not showcase these recovered values until after their environmental temperatures return to non-extreme conditions, and likely their endosymbiont populations, who are capable of providing corals with 100% of their daily energy requirements, are fully returned (Grottoli et al. 2006). Prior to the knowledge of extreme bleaching preceding collection of the data presented here, energy reserves of surviving corals five years after transplantation were expected to mirror results seen 17-months after transplantation: higher concentrations in corals placed in the nearshore environment, supporting evidence of their successful acclimatization from the offshore environment. In reality, the lack of difference in bulk protein and lipid concentrations between coral species and treatments does not support the acclimatization potential of these corals but instead may provide insight into the bleaching response of these species.

For proteins, these results nearly mirror those seen 17 months after transplantation, besides a decrease in protein concentration of *P. strigosa* native to the offshore between 9 and 17 months (Baumann et al. 2021). Previous work with Caribbean coral (*Montastraea franksi*) is supportive of relatively unchanged pooled concentrations of glycerol, protein, and tissue biomass when comparing pre- and post-bleached levels (Edmunds et al. 2003). However, an experiment subjecting the same species studied here to elevated temperature for 95 days was found to significantly reduce the protein concentrations in *S. siderea* and *P. strigosa* fragments from both nearshore and backshore reefs in Belize (Aichelman et al. 2021). The difference in protein concentrations observed by Aichelman et al. (2021) could be associated with testing these corals in a lab-based experimental setting, where they had limited access to nutrition levels, whereas the corals in our field study saw natural abundances of particulate and dissolved nutrition sources *in-situ* every day, which may have prevented a decrease in protein concentration.

Lipids have the highest caloric content when compared to proteins and carbohydrates and serve as longer-term energy reserves as opposed to short-term carbohydrates (Gnaiger and Bitterlich, 1984; Grottoli et al., 2004). Lipids are an important source of energy in the reproduction of corals, and high lipid stores are known to assist in recovery from and resistance to bleaching (Anthony et al., 2009; Ward, 1995). In many coral species, lipids appear to be the first line of defense and energy source for corals under bleaching conditions. The lipid profiles of healthy, partially bleached, and fully bleached corals have been found to be significantly different (Bachok et al. 2006). In a study looking at the recovery of multiple tropical corals following recurring bleaching events, the researchers believed that the reason one species (*P. astreoides*) was unable to successfully recover within the span of 11 months was due to its lower concentrations of lipid reserves compared to the species which successfully recovered (Schoepf et al., 2015). As a result of lower baseline lipid storage, it is likely that these species had to use up its protein and carbohydrate stores to endure during bleaching and therefore was unable to achieve full recovery within 11 months (Schoepf et al., 2015). Given that corals at both sites in this experiment were subjected to extreme temperatures prior to fragment collection and lab-based processing, it is not surprising that both species displayed low levels of bulk lipids.

While proteins and lipids did not differ between species or transplant treatments, there was a clear species-specific difference in the carbohydrate concentration where *P. strigosa* displayed significantly lower levels than *S. siderea* five years after transplantation. While this species-specific difference in carbohydrates was not displayed by these same corals at earlier time points in this experiment (Baumann et al. 2021), previous work has shown general trends of *S. siderea* maintaining higher concentrations of host carbohydrates as compared to *P. strigosa* (Bove et al. 2022a). Carbohydrate reserves are also known to be impacted under bleaching stress. Of the three macromolecules assessed here, carbohydrates provide the least amount of energy to coral hosts (Anthony et al. 2002). Carbohydrates can be rapidly broken down for energy as compared to protein or lipid molecules when considering enthalpies of combustion (Grottoli et al. 2004), and have been found to rapidly deplete under thermal stress (Kochman et al. 2021). Therefore, the lower concentrations of carbohydrates in *P. strigosa* may be indicative of higher bleaching impacts felt by this species compared to *S. siderea*. Carbohydrates can also be acquired via heterotrophy of the coral host aside from carbohydrates deposited by photosynthetic processing by endosymbionts, potentially pointing to increased heterotrophic capacity of *S. siderea* compared to *P. strigosa* (Dellisanti et al. 2023). While bleaching conditions were experienced at both sites, the lower bleaching tolerance in *P. strigosa* would cause these corals to more heavily deplete their storages of carbohydrates, mirroring the results of *P. strigosa* displaying lower carbohydrate concentrations than *S. siderea* across all transplant treatments. Additionally, *P. strigosa* native to the offshore had significantly higher carbohydrate concentrations compared to all other treatments. However, considering that the average amount of carbohydrates found in this group were ∼0.02 mg/mg, these values suggest that the carbohydrate reserves of these corals were still very much impacted by bleaching stress. The order that energy reserves are recovered following stress events including bleaching is known to differ by species (Rodrigues and Grottoli 2007). Given carbohydrates’ ability to be rapidly synthesized as a form of small, quick stored energy (Kochman et al. 2021), it is possible that *P. strigosa* in the offshore site were able to start restocking these reserves first as thermal stress began to subside at the offshore site. The fact that photosynthetic efficiency values of *P. strigosa* placed in the offshore were elevated compared to *P. strigosa* placed in the nearshore supports the feasibility of this scenario.

The results presented here – especially of survival, growth, and bulk carbohydrate concentrations — offer evidence to support that offshore *S. siderea* transplanted to the nearshore have high acclimatization potential compared to all other species and treatment groups. Meanwhile growth and mortality data convey that offshore *P. strigosa* and nearshore populations of both species appear to be limited in their acclimatization potential, potentially due to local adaptation. The displayed species-difference in carbohydrate concentrations may reflect species-specific recovery and or bleaching patterns. Surface level differences in the acclimatization potential of *P. strigosa* and *S. siderea* could be explained by general differences in the metabolic rates and enzyme activities of separate species (Seebacher et al. 2015). However, at the same time, there are questions about whether acclimatization of populations can be developed on a scale that meets the pace of the changing climate. Even in the case of corals acquiring increased thermal tolerance, the fact that populations, including all those in this study, still exhibit high mortality as a result of bleaching is evidence that this tolerance mechanism is not taking place fast enough to prepare for more extreme stressor events in the future (Coles et al. 2018).

## Conclusion

To summarize, offshore *S. siderea* was the only group that continued to display any evidence of acclimatization five years after transplantation, while the groups of *P. strigosa* seem to be limited in their acclimatization due to local adaptation to their native sites and sensitivity to bleaching. The implications of these results are relevant to restoration and outplanting efforts. Corals that exhibit high acclimatization potential can be successful options for coral outplanting efforts. On the other hand, corals that appear to be relying on local adaptation potential as a tolerance mechanism should be kept in their native environment with restoration efforts. *Siderastrea. siderea* may be a useful target species when the focus of these efforts is within nearshore environments whereas *P. strigosa* may be more useful when being maintained within or near their native environments

## Limitations

It is important to note the three primary limitations in this work. Firstly, stony coral tissue loss disease (SCTLD), has severely impacted Caribbean corals through a relatively unconfirmed mechanistic basis (Walton et al. 2018). While these two species display stress tolerant characteristics, they are both known to be particularly susceptible to coral diseases, including Stony Coral Tissue Loss Disease (Alvarez-Filip et al. 2022). Recorded mortality from SCTLD at the offshore site in this study was reported by the Fragments of Hope team in mid-April of last year (2023), meaning that this disease likely arrived in the winter of 2022–2023. The presence of SCTLD here is a confounding factor especially for the mortality and growth data of corals placed at the offshore site. However there was no evidence of SCTLD presence at the NS site. While there was some evidence of SCTLD on the remaining samples when we arrived in the field, most of the living colonies seemed relatively unscathed from the disease. Apart from total recorded mortality from the three previous time points (*P. strigosa* n = 3; *S. siderea* n = 26), it is possible that corals mechanically detached from their transplant table as a result of high wave energy or storm action. Secondly, the extreme global heatwave that occurred in the summer of 2023 causing massive bleaching was an unexpected event in the planning of this project (Birkart and Álvarez-Filip 2025). Nonetheless, given that all the corals at each site experienced the same extreme temperatures, this event functioned as a stress-test and allowed for an investigation of differentiated resiliency between species and location after five years of acclimatization by transplant site. Finally, the limited number of site visits over the timeline of this work presents a constrained discussion of results. Again, the time points of these visits were at 0, 3, 9, 17, and 60 months following transplantation. There is good coverage of the samples and their well-being in the short term (0-17 months) but a large gap between the last two time points.

## Acknowledgements

This study was made possible by Belize Fund for a Sustainable Future research grant to Fragments of Hope^Ltd^, an Amherst College Gregory S. Call Research Grant to CH, a Mount Holyoke College Faculty Development Grant to JHB, startup funds from Bates College to JHB, and startup funds from Ursinus College to CBB.

## Permitting

The fieldwork was done under a marine research permit granted by the Belize Fisheries Department to Fragments of Hope^Ltd^ and collaborators. Transport of corals was conducted following Belize, US, and international regulations with permits from the Belize Fisheries Department, US Fish, and Wildlife Services, and the Convention on International Trade of Endangered Species (CITES).

