## Supplemental Figure and Tables for "Acclimatization potential in stress tolerant Caribbean corals - outcomes of a 5 year cross-shelf reciprocal transplant experiment"

### Supplemental files

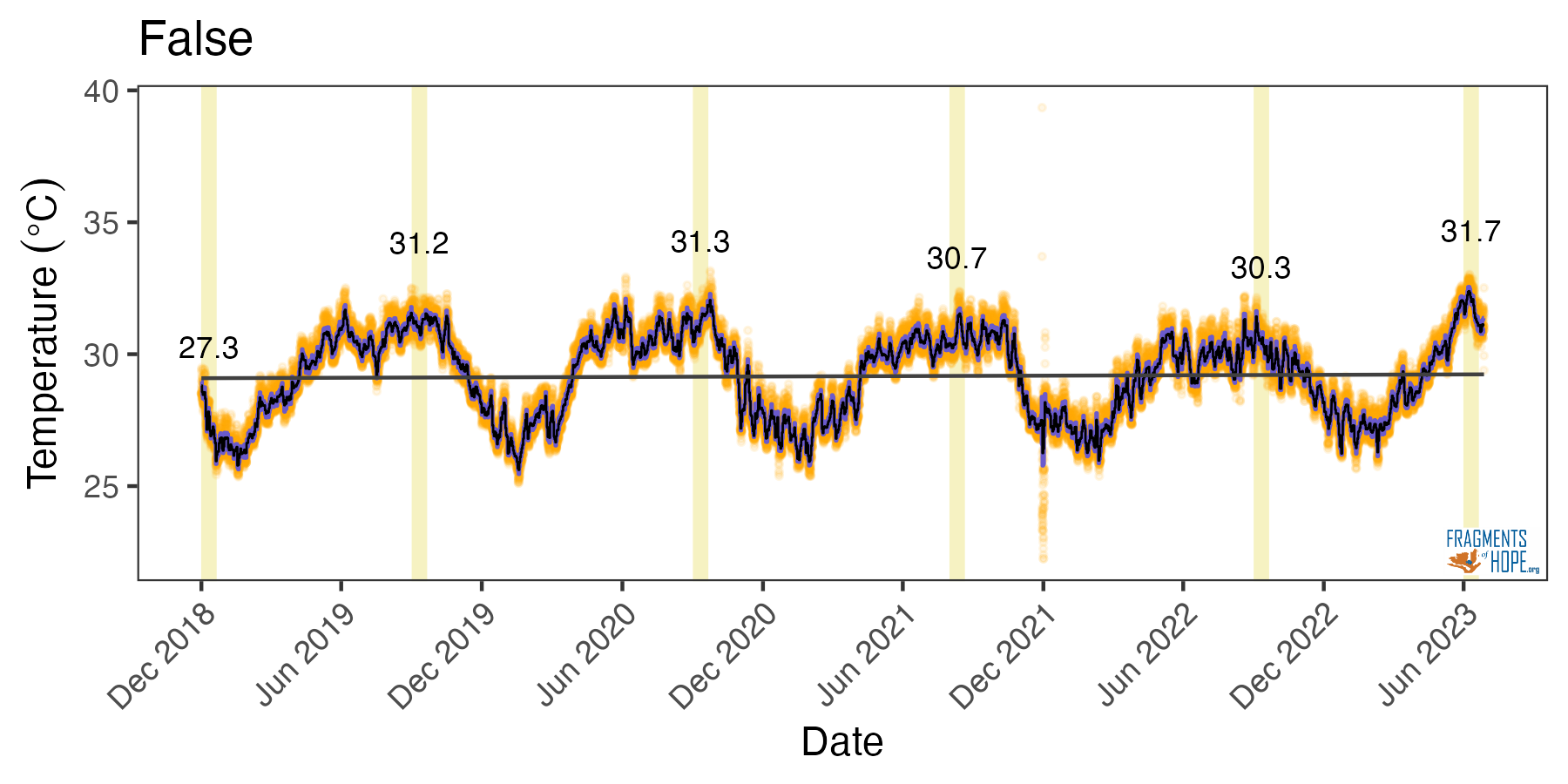

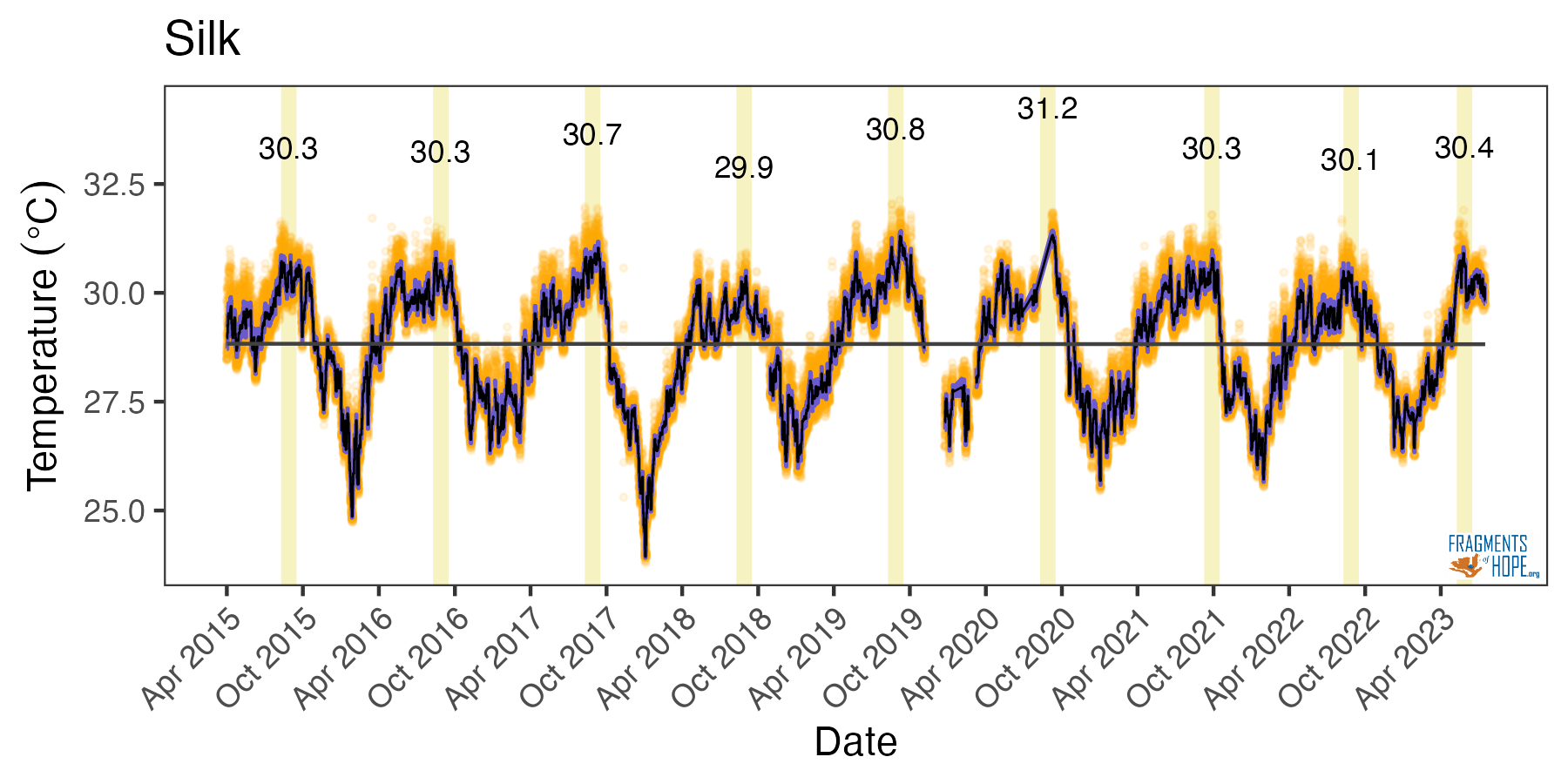

**17-month time point**

**A**

**B**

**Fig S1:** In-situ temperature at each site across the 60 month study. A.) Nearshore site (False Caye), B.) Offshore site (Silk Caye). The numbers above the peaks are the maximum average daily temperatures for each year. The red dashed lines approximate the 17-month time point of the experiment, with the ~0.4°C difference in maximum average daily *in-situ* temperature recorded by HOBO^®^ V2 data loggers.

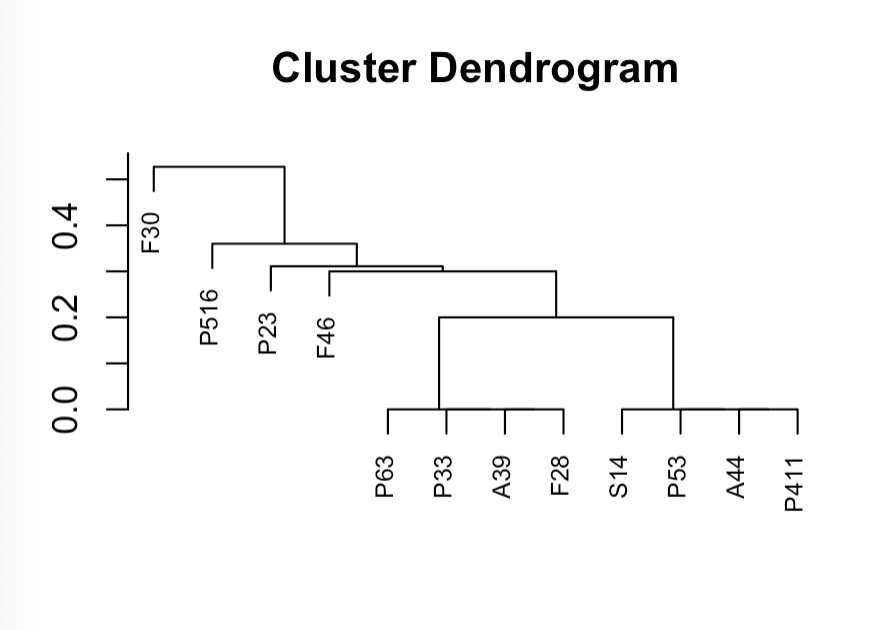

Height

**Figure S2:** Dendrogram created with 2bRADseq pipeline for *P. strigosa* samples.

**Figure S3:** Dendrogram created with 2bRADseq pipeline for *S. siderea* samples.

| **Species** | **Temporary ID** | **Transplant Reef** | **2b-RAD ID successful?** | **Home Reef** | **Treatment** |
| --- | --- | --- | --- | --- | --- |
| *P. strigosa* | False30 | NS | No | Unknown | Unknown |
| *P. strigosa* | False46 | NS | No | Unknown | Unknown |
| *P. strigosa* | False28 | NS | No | Unknown | Unknown |
| *P. strigosa* | Silk14 | OS | No | Unknown | Unknown |
| *S. siderea* | False7 | NS | No | Unknown | Unknown |
| *S. siderea* | False8 | NS | No | Unknown | Unknown |
| *S. siderea* | False 12 | NS | Yes | OS | Transplant to NS |
| *S. siderea* | False18 | NS | Yes | NS | NS Native |
| *S. siderea* | False19 | NS | No | Unknown | Unknown |
| *S. siderea* | False23 | NS | No | Unknown | Unknown |
| *S. siderea* | False3 | NS | Yes | NS | NS Native |
| *S. siderea* | False20 | NS | No | Unknown | Unknown |
| *S. siderea* | Silk23 | OS | No | Unknown | Unknown |

**Table S1:** Coral fragments with degraded labels 60 months after transplantation, including their location (transplant reef) and whether or not identification by 2b-RAD was successful.

|  | Calcification | Fv/Fm | Protein | Lipid | Carbs | Chl a | Chl c |
| --- | --- | --- | --- | --- | --- | --- | --- |
| Intercept (NS Native) | 3.473** | 0.548*** | 0.207*** | 0.114** | 0.021*** | 0.501 | 2.326* |
|  | (1.217) | (0.016) | (0.022) | (0.033) | (0.004) | (0.357) | (1.054) |
| OS Native | -0.781 | 0.009 | -0.089* | -0.002 | 0.012* | 0.663 | 1.737 |
|  | (0.784) | (0.021) | (0.029) | (0.045) | (0.006) | (0.483) | (1.427) |
| Transplant to NS | 3.303*** | 0.033 | 0.000 | -0.059 | 0.001 | 0.993+ | 2.766 |
|  | (0.778) | (0.021) | (0.032) | (0.049) | (0.006) | (0.535) | (1.580) |
| SD (Intercept colony_ID) | 2.675 | 0.009 | 0.003 |  |  |  |  |
| SD (Observations) | 1.378 | 0.046 | 0.048 |  |  |  |  |
| Num.Obs. | 27 | 30 | 15 | 15 | 15 | 15 | 15 |
| R2 |  |  |  | 0.130 | 0.330 | 0.237 | 0.212 |
| R2 Adj. |  |  |  | -0.015 | 0.218 | 0.109 | 0.081 |
| R2 Marg. | 0.273 | 0.084 | 0.467 |  |  |  |  |
| R2 Cond. | 0.848 | 0.119 |  |  |  |  |  |
| F |  |  |  | 0.899 | 2.950 | 1.860 | 1.615 |

**Table S2:** Model coefficients for all physiological parameters measured in *S. siderea*. Values represent model coefficient estimates with standard errors in parentheses. The intercept represents the NS Native treatment group (reference level), and remaining coefficients reflect differences relative to this group. Linear mixed-effects models with colony as a random effect were used for calcification, Fv/Fm, and protein, for which marginal and conditional R² values are reported; simple linear models were used for lipid, carbohydrate, chlorophyll a, and chlorophyll c2 where LMMs produced singular fits, for which R² and adjusted R² values are reported.

| Parameter | Contrast | Estimate | SE | df | p-value |  |
| --- | --- | --- | --- | --- | --- | --- |
| Calcification | NS Native - OS Native | 0.781 | 0.801 | 20.748 | 0.600 |  |
|  | NS Native - Transplant to NS | -3.303 | 0.792 | 20.540 | 0.001 | ** |
|  | OS Native - Transplant to NS | -4.084 | 0.654 | 19.249 | 0.000 | *** |
| Fv/Fm | NS Native - OS Native | -0.009 | 0.022 | 26.999 | 0.914 |  |
|  | NS Native - Transplant to NS | -0.033 | 0.022 | 26.728 | 0.297 |  |
|  | OS Native - Transplant to NS | -0.024 | 0.021 | 25.336 | 0.477 |  |
| Protein | NS Native - OS Native | 0.089 | 0.032 | 11.487 | 0.045 | * |
|  | NS Native - Transplant to NS | 0.000 | 0.034 | 10.097 | 1.000 |  |
|  | OS Native - Transplant to NS | -0.089 | 0.032 | 8.911 | 0.049 | * |
| Lipid | NS Native - OS Native | 0.002 | 0.045 | 12.000 | 0.999 |  |
|  | NS Native - Transplant to NS | 0.059 | 0.049 | 12.000 | 0.483 |  |
|  | OS Native - Transplant to NS | 0.057 | 0.047 | 12.000 | 0.479 |  |
| Carbs | NS Native - OS Native | -0.012 | 0.006 | 12.000 | 0.109 |  |
|  | NS Native - Transplant to NS | -0.001 | 0.006 | 12.000 | 0.973 |  |
|  | OS Native - Transplant to NS | 0.011 | 0.006 | 12.000 | 0.197 |  |
| Chl a | NS Native - OS Native | -0.663 | 0.483 | 12.000 | 0.385 |  |
|  | NS Native - Transplant to NS | -0.993 | 0.535 | 12.000 | 0.194 |  |
|  | OS Native - Transplant to NS | -0.330 | 0.515 | 12.000 | 0.801 |  |
| Chl c | NS Native - OS Native | -1.737 | 1.427 | 12.000 | 0.466 |  |
|  | NS Native - Transplant to NS | -2.766 | 1.580 | 12.000 | 0.227 |  |
|  | OS Native - Transplant to NS | -1.029 | 1.521 | 12.000 | 0.781 |  |

**Table S3:** Pairwise contrasts from estimated marginal means (EMMs) for all physiological parameters measured in *S. siderea*. Estimates represent the difference between group means, with standard error (SE), degrees of freedom (df), and Tukey-adjusted p-values reported for each contrast. Significance codes: * *p* < 0.05, ** *p* < 0.01, *** *p* < 0.001.

|  | Fv/Fm | Calcification | Protein | Lipid | Carbs | Chl a | Chl c |
| --- | --- | --- | --- | --- | --- | --- | --- |
| Intercept (NS Native) | 0.467*** | 0.149*** | 0.149*** | 0.062+ | 0.007*** | 0.261 | 1.646* |
|  | (0.012) | (0.024) | (0.024) | (0.029) | (0.001) | (0.180) | (0.521) |
| OS Native | 0.083** | -0.118* | -0.118* | 0.060 | 0.013*** | 0.342 | 0.795 |
|  | (0.023) | (0.040) | (0.040) | (0.048) | (0.002) | (0.293) | (0.850) |
| Transplant to OS | 0.111*** | -0.033 | -0.033 | 0.056 | 0.003 | 1.733*** | 5.406*** |
|  | (0.027) | (0.037) | (0.037) | (0.044) | (0.002) | (0.269) | (0.781) |
| SD (Intercept colony_ID) | 0.008 |  |  |  |  |  |  |
| SD (Observations) | 0.047 |  |  |  |  |  |  |
| Num.Obs. | 26 | 12 | 12 | 12 | 12 | 12 | 12 |
| R2 |  | 0.498 | 0.498 | 0.204 | 0.772 | 0.830 | 0.853 |
| R2 Adj. |  | 0.387 | 0.387 | 0.027 | 0.721 | 0.792 | 0.820 |
| R2 Marg. | 0.498 |  |  |  |  |  |  |
| R2 Cond. | 0.512 |  |  |  |  |  |  |
| F |  | 4.466 | 4.466 | 1.154 | 15.222 | 21.986 | 26.110 |

**Table S4:** Model coefficients for all physiological parameters measured in *P. strigosa*. Values represent model coefficient estimates with standard errors in parentheses. The intercept represents the NS Native treatment group (reference level), and remaining coefficients reflect differences relative to this group. A linear mixed-effects model with colony as a random effect was used for Fv/Fm, for which marginal and conditional R² values are reported; simple linear models were used for calcification, protein, lipid, carbohydrate, chlorophyll a, and chlorophyll c2 where LMMs produced singular fits, for which R² and adjusted R² values are reported.

| Parameter | Contrast | Estimate | SE | df | p-value |  |
| --- | --- | --- | --- | --- | --- | --- |
| Fv/Fm | NS Native - OS Native | -0.083 | 0.025 | 22.961 | 0.007 | ** |
|  | NS Native - Transplant to OS | -0.111 | 0.028 | 22.210 | 0.002 | ** |
|  | OS Native - Transplant to OS | -0.028 | 0.034 | 22.979 | 0.696 |  |
| Calcification | NS Native - OS Native | 0.118 | 0.040 | 9.000 | 0.038 | * |
|  | NS Native - Transplant to OS | 0.033 | 0.037 | 9.000 | 0.649 |  |
|  | OS Native - Transplant to OS | -0.085 | 0.042 | 9.000 | 0.158 |  |
| Protein | NS Native - OS Native | 0.118 | 0.040 | 9.000 | 0.038 | * |
|  | NS Native - Transplant to OS | 0.033 | 0.037 | 9.000 | 0.649 |  |
|  | OS Native - Transplant to OS | -0.085 | 0.042 | 9.000 | 0.158 |  |
| Lipid | NS Native - OS Native | -0.060 | 0.048 | 9.000 | 0.450 |  |
|  | NS Native - Transplant to OS | -0.056 | 0.044 | 9.000 | 0.436 |  |
|  | OS Native - Transplant to OS | 0.004 | 0.050 | 9.000 | 0.997 |  |
| Carbs | NS Native - OS Native | -0.013 | 0.002 | 9.000 | 0.001 | ** |
|  | NS Native - Transplant to OS | -0.003 | 0.002 | 9.000 | 0.339 |  |
|  | OS Native - Transplant to OS | 0.010 | 0.002 | 9.000 | 0.009 | ** |
| Chl a | NS Native - OS Native | -0.342 | 0.293 | 9.000 | 0.501 |  |
|  | NS Native - Transplant to OS | -1.733 | 0.269 | 9.000 | 0.000 | *** |
|  | OS Native - Transplant to OS | -1.391 | 0.307 | 9.000 | 0.004 | ** |
| Chl c | NS Native - OS Native | -0.795 | 0.850 | 9.000 | 0.633 |  |
|  | NS Native - Transplant to OS | -5.406 | 0.781 | 9.000 | 0.000 | *** |
|  | OS Native - Transplant to OS | -4.611 | 0.889 | 9.000 | 0.001 | ** |

**Table S5:** Pairwise contrasts from estimated marginal means (EMMs) for all physiological parameters measured in *P. strigosa*. Estimates represent the difference between group means, with standard error (SE), degrees of freedom (df), and Tukey-adjusted p-values reported for each contrast. Significance codes: * *p* < 0.05, ** *p* < 0.01, *** *p* < 0.001.
